# Global Neural Oscillations Underlie Performance Variability and Attentional State Fluctuations in Humans

**DOI:** 10.64898/2026.03.31.715029

**Authors:** Joaquín Herrero, Rodrigo Henríquez-Ch, Alejandra Figueroa-Vargas, Reinaldo Uribe-San Martin, Christian Cantillano, Pablo Fuentealba, Patricio Mellado, Jaime Godoy, Pablo Billeke, Francisco Aboitiz

## Abstract

Fluctuations in attentional states, such as mind-wandering (MW), are associated with critical variability in task performance. While fMRI studies highlight the opposing roles of task-positive (e.g., dorsal attention network) and task-negative (e.g., default mode network) systems, the electrophysiological mechanisms underlying these dynamics remain poorly understood. Using intracranial electrocorticography in humans performing a sustained attention task, we identified global oscillatory dynamics linked to attentional shifts. MW was characterized by (i) reduced theta (θ) and alpha (□) power, (ii) decreased aperiodic signal components, indicating a shift toward cortical inhibition, (iii) enhanced phase synchronization across networks, and (iv) strengthened θ phase-behavior correlations (ρ). These features support a non-network-specific framework in which low-frequency θ dynamics—captured by both θ power and ρ—are associated with attentional fluctuations, while aperiodic offset relates to attentional state indirectly through its association with ρ (Structural Equation Modeling: power→state β = −0.118, p = 0.002; ρ→state β = 0.246, p < 0.001; offset→ρ β = −0.222, p < 0.001). Our study provides a unified neurophysiological framework for understanding how spontaneous neural activity can drive attentional fluctuations and performance variability, with implications for research on attention, learning, and neuropsychiatric disorders.

## Introduction

To navigate a multi-demanding environment, we must flexibly focus on the task at hand while remaining alert to other needs and concurrent internal processes. For instance, while writing a report at work, our attention may be interrupted by internal thoughts, such as wondering whether we locked the door at home or remembering that our son has upcoming tasks at school. Fluctuations in attentional states are often associated with shifts in immediate internal focus, impacting task performance, where moments of distraction or reduced focus lead to increased variability in outcomes. While this process could be a key physiological function[1], it may also lead to weakness in crucial tasks, potentially resulting in serious consequences—for example, a pilot combating an extended forest fire during a long shift. A salient manifestation of these fluctuations is called mind-wandering (MW), which refers to the spontaneous shift of attention away from the task at hand. Crucially, rather than being random, these shifts are thought to be governed by neural mechanisms that dynamically regulate attentional focus, interplaying between endogenous cognitive processes and external task demands[2,3].

A candidate neural mechanism responsible for such attentional fluctuations is ongoing oscillatory brain activity. Emerging evidence suggests that neural oscillations play a fundamental role in the emergence of distinct attentional states [4–7]. Indeed, spontaneous activity patterns are related to the performance of cognitive processes and changes in learning (e.g., in visual, sensory-motor, or other processes) and are implicated in behavioral abnormalities across various clinical conditions [8–18].

Evidence of neuronal dynamics underlying attentional shift primarily comes from slow changes in metabolic activity, which shape brain networks with distinct activity patterns. On the one hand, task-positive networks, which include the Dorsal Attention Network (DAN), Ventral Attention Network (VAN), and Fronto-Parietal Control (FPC) Network, support goal-oriented, externally focused cognitive processes [19–22]. On the other hand, task-negative networks, primarily the Default Mode Network (DMN), are active during internally focused thinking and deactivate during externally directed tasks [23]. Such an antagonistic relationship governs the interaction between these networks: sustained attention is marked by heightened task-positive activity alongside suppression of task-negative regions, whereas attentional lapses, including MW, are characterized by opposite patterns. It is proposed that this dynamic balance is essential for cognitive adaptive cognitive control [20,24].

Despite extensive research on slow fluctuations in metabolic activity, the electrophysiological mechanisms underlying transitions between different attentional states are poorly understood. Low-frequency brain oscillations, particularly theta oscillations (4–8 Hz), have been identified as a key mechanism in several cognitive processes [7,17,25–36], including attention modulation [4,5,7,37,38]. These oscillations play a critical role in predicting sustained attention performance, regardless of task and context[39,40]. Notably, brain networks involved in task performance display distinct oscillatory patterns compared to those less engaged, contributing to variations in reaction times and attentional focus[41]. However, despite the recognized importance of these oscillatory mechanisms, the specific dynamics driving performance variability across different attentional states remain largely unexplored.

Another key physiological feature linked to attentional states is the excitatory/inhibitory (E/I) balance, which can be measured through the aperiodic component of the electrical brain signal [42–44]. While periodic activity—categorized into bands such as theta—has been widely studied for its role in perceptual, cognitive, and motor processes, aperiodic components offer critical insights into synaptic balance [44]. Aperiodic measures, such as the slope and offset of the power spectral density, also vary across brain states. For example, a decreased slope during sleep and anesthesia reflects increased cortical inhibitory gain [45,46]. Moreover, aperiodic components are linked to cognitive and perceptual processes. For instance, attentional processes with varying sensory demands have been differentiated by aperiodic components [47], highlighting their relevance in capturing the inherent fluctuations in attentional states. These findings underscore the importance of aperiodic activity in understanding the dynamic nature of attention and its underlying neural mechanisms.

Despite accumulated evidence, the precise neurophysiological mechanisms underlying attentional fluctuations remain elusive. This gap impedes our understanding of the physiological principles governing attentional shifts and the development of strategies to maintain focus under high external demands. To address this, we leveraged the high spatiotemporal precision of electrocorticography (ECoG) in nine patients with pharmacoresistant epilepsy during a sustained attention task. This approach allowed us to simultaneously evaluate oscillatory dynamics and aperiodic components across task-positive (dorsal attention, ventral attention, frontoparietal control) and task-negative (default mode) networks.

We hypothesized that modulations in low-frequency oscillations—specifically theta and alpha power and phase—underlie the transitions between sustained attention and off-task states, such as mind-wandering. Our results identify a global, non-network-specific mechanism in which reductions in theta/alpha power and phase jointly modulate performance variability. Notably, these effects were accompanied by a generalized shift in aperiodic components across the cortex, suggesting a widespread modulation of the excitatory-inhibitory balance during mind-wandering. By demonstrating that these dynamics occur across the entire cortex rather than being localized to specific systems, our findings challenge prevailing models that attribute mind-wandering exclusively to the dominance of the default mode network.

## Results

### Behavioral variability differences in attentional states

We recorded ECoG from nine pharmaco-resistant epilepsy patients (mean age: 27 years, SD: 9.6; 7 female) while they performed a Sustained Attention Response Task (SART; Fig. 1a). In this task, participants had to determine whether the visual stimuli presented on the screen were targets or non-targets. The target frequency was low, and the inter-stimulus interval was long (see Methods for details) to induce fluctuations in attentional focus. After 5-8 trials, a question appeared on the screen to collect participants’ subjective reports on their attentional focus. The Thought Sampling Questions (TSQ) were used to classify attentional states into three distinct categories: “On-Task” (ON), “Off-Task” (OFF), and “Sleep” (SL). The electrodes for all subjects were independently positioned within the MNI space and subsequently distributed across the brain networks of interest (see Method, Fig. 1b).

**Figure 1:**
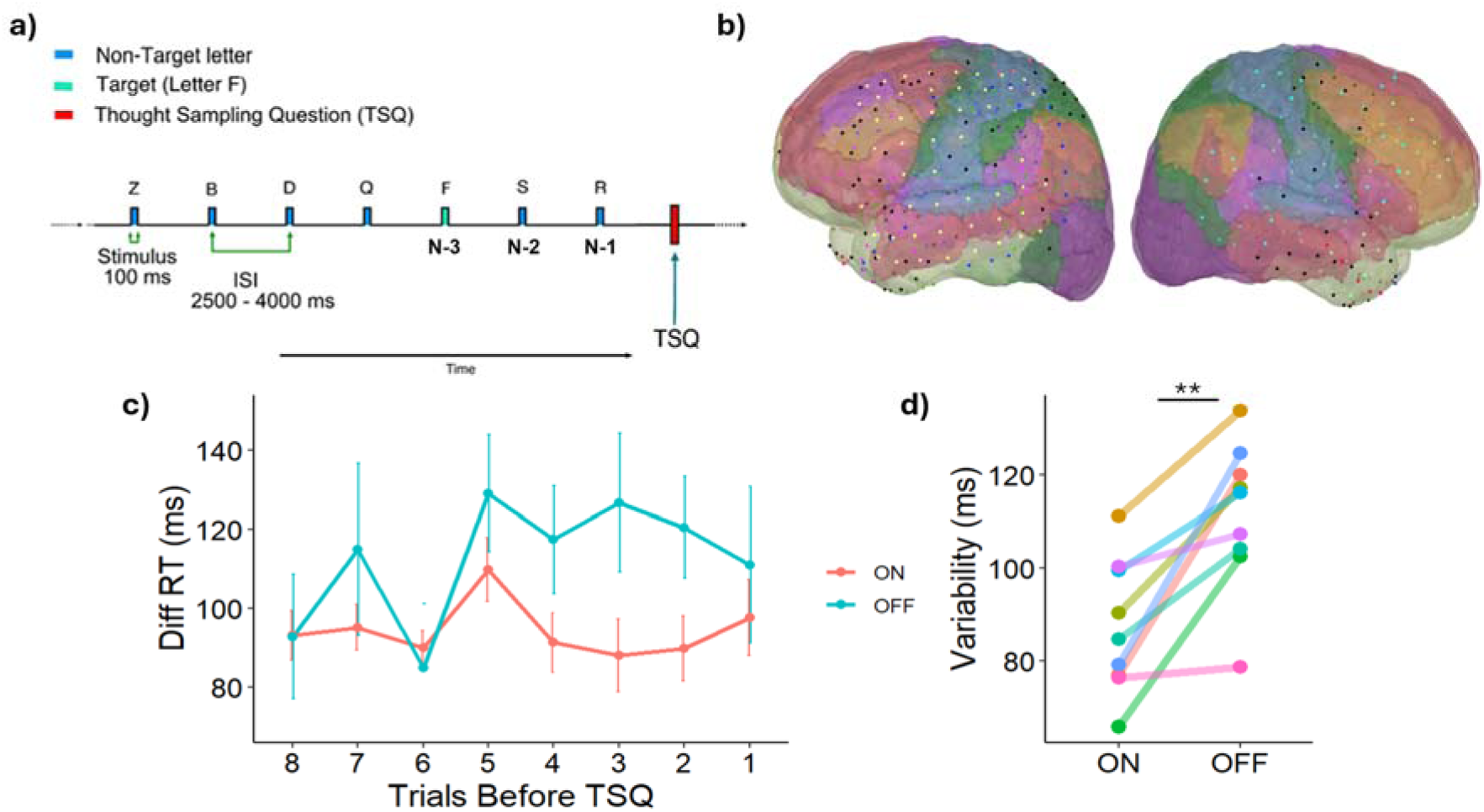
Task, Electrode Localization, and Task Performance. **(a)** Sustained Attention Response Task (SART). The target stimulus is the letter “F” (infrequent). After 5 or 8 trials (letters), a Thought Sampling Question (TSQ) with a forced-choice response is presented. Trials preceding the TSQ are labeled as N-x. **(b)** Electrode placements from nine participants represented in MNI space. Each color corresponds to a different participant, with black points indicating epileptogenic electrodes. The background color represents functional brain networks, with the networks of interest displayed as follows: Default Mode Network (DMN, red), Dorsal Attention Network (DAN, purple), Ventral Attention Network (VAN, green), and Frontoparietal Control Network (FPC, orange). **(c)** Trial-to-trial differences in reaction times (ms) for the eight trials preceding a TSQ, categorized by ON (red line) and OFF (blue line) states. **(d)** Reaction time variability across the four trials preceding each TSQ response. Each color represents a patient. (\*\**p* < .01).

We first analyzed hit rates for both target and non-target trials to assess task performance. The hit rate for non-target trials was 99.1% ± 1.3%, whereas for target trials, it was significantly lower at 85.7% ± 6.3% (W = .1, p = .0004). RT also differed significantly between trial types: non-target trials had a mean RT of 471.1 ± 39.1 ms, while target trials were associated with a slower RT of 602.6 ± 60.2 ms (W = 80, p = .0008). These findings are consistent with previous evidence indicating that participants maintain high accuracy in low-complexity tasks [48]. Furthermore, the low frequency of target trials contributed to prolonged RTs, likely reflecting an expectancy bias toward non-target stimuli [7,34]. Overall, these results confirm that the task was effectively implemented, eliciting distinct performance patterns between target and non-target trials and reinforcing the cognitive demands of sustained attention over extended periods.

To characterize the temporal dynamics of MW onset, we analyzed RTs in trials preceding the presentation of the TSQs. Across attentional states, no significant differences in RTs were observed. However, previous studies have consistently linked RT variability to MW states[48–52]. To examine trial-to-trial fluctuations in RT, we computed the difference in RT between adjacent trials as a measure of variability (Fig. 1c). When assessed cumulatively variability across the four trials preceding the TSQ, OFF states exhibited significantly greater RT variability compared to both ON states (t(14) = -2.4, p =.03) (Fig. 1d). In contrast, when analyzing all preceding trials collectively, no significant differences in RT variability were observed (t(13) = -1.9, p = .07). A Bayesian regression model further supported these findings, identifying cumulative RT variability across the last four trials and variability in the most recent trial as the best predictors of MW dynamics (β1(median) = -0.65, CI [-1.1 -0.8]; DIC = 316). These results underscore the relevance of RT variability as a marker of MW onset and highlight the complex interplay between attentional states and behavioral fluctuations.

In summary, the behavioral findings demonstrate expected task performance, marked by a lower hit rate and prolonged reaction times during target trials compared to non-target trials. These differences likely reflect distinct cognitive processes related to the novelty or salience of target stimuli. Notably, the OFF state exhibited sustained variability across trials preceding the TSQ, with high variability in the final 4 trials (corresponding to 10–14 seconds before the MW report) emerging as a key marker of attentional state transitions. This pattern highlights the gradual temporal dynamics underlying internal and external attention shifts.

### Low-frequencies predominate in OFF states across Brain Networks of Interest

We investigated neural oscillatory dynamics by comparing ON and OFF attentional states. The electrodes included in these analyses were classified into one of three network categories—task-positive (VAN, DAN, FPC), task-negative (DMN), or other networks (SM, limbic). Time–frequency analyses were performed time-locked to the onset of non-target stimuli. Additionally, we considered the periods preceding stimulus presentation (negative time windows) to distinguish patterns of spontaneous neural activity. To enable network-level comparisons, we focused on three functionally defined networks: the task-negative network (TN), the task-positive network (TP), and other networks (OTHERS, see Methods for further details). Given the variability in intra-network oscillatory activity[19,21] and the differences in anatomical localization within and across networks (Table 3), we applied a linear mixed-effects model (LMM). The attentional state (ON/OFF) was included as a fixed effect, while random effects accounted for electrode-specific variability.

Our analysis revealed significant differences in low-frequency oscillatory power across the networks of interest. Specifically, the task-negative network exhibited a pronounced reduction in power during the periods preceding stimulus presentation, particularly in the theta band (Theta [4-8 Hz, -1.5 - 0.8 Sec]; β = -135.9; t = -4.3; Alpha [8-12 Hz, -1.5 - 0.8 Sec]; β = -33.8, t = -3.1; sea also Fig. 2a). Similarly, the task-positive and other networks showed comparable dynamics, with power reductions primarily observed in the theta (Theta TP β = -98.4, t = -2.8; Theta Others β = -120.2, t = -3.5) and alpha bands (Alpha TP β = - 18.7, t = -1.7; Alpha Others β = -50.9, t = -3.0, see Fig. 2b–c). When considering all electrodes together (Fig. 2d), this effect was more pronounced and closely resembled the pattern observed in the task-negative network, with significant decreases in both theta and alpha spectral power during the OFF state (Theta β = -118.6, t = -5.9; Alpha β = -35.2, t = - 4.2). Accordingly, when examining the distribution of this differential activity across the networks of interest (Fig. 2e), we found that theta band differences occurring between -1 second and -0.1 seconds relative to the event were uniformly distributed across networks (*X²*(3,379) = 1.3, *p* = 0.5). These findings highlight distinct oscillatory signatures associated with attentional states, suggesting that global dynamics govern attentional fluctuations.

**Figure 2:**
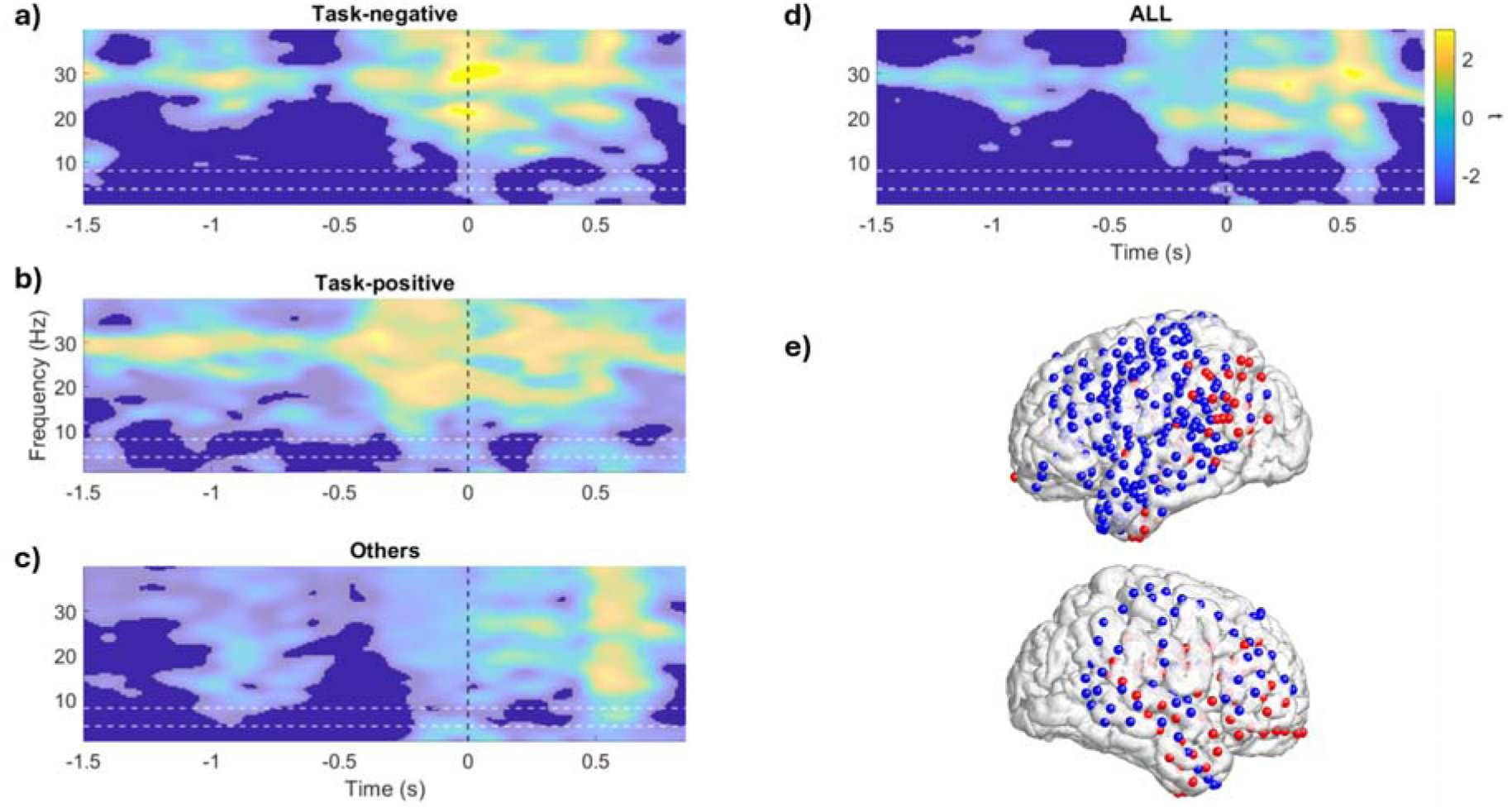
Differences Between ON and OFF States Across Brain Networks of Interest. **(a)** Time–frequency decomposition (TFD) for task-negative electrodes is shown for both the periods preceding stimulus presentation (negative x-axis) and the period following stimulus presentation (positive x-axis). Zero represents letter presentation (trial onset). **(b–d)** TFD for task-positive networks (b), other networks (c), and all networks combined (d), respectively. (a-d) Color represents the t-value of the LMM for the attentional state regressor. The highlighted area denotes statistically significant time-frequency differences, identified using a false discovery rate (FDR) threshold of q<0.05. **(e)** Electrode distribution illustrating the effects presented in panel (d). Electrodes shown in blue contribute to the global effect observed in (d), while electrodes in red do not contribute to the global effect.

### Inhibitory mechanisms predominate in OFF states

To elucidate changes associated with the I/E balance in brain networks involved in MW, we examined the aperiodic components of spectral power density. Shifts in aperiodic components, specifically in the exponent (slope) and intercept (offset), are thought to indirectly reflect synaptic-level alterations in I/E balance [42,44]. To assess these dynamics, we applied an LMM to analyze aperiodic components within each brain network of interest.

Our analysis revealed decreases in both the slope and offset during OFF states when all electrodes were considered collectively (Fig. 3a). A similar pattern emerged when electrodes were grouped into the TN, TP, and other networks, except for the slope in the TN (β = -0.04, t = -1.14, C.I. [-0.09, 0.02]) and TP networks (β = -0.05, t = -1.6, C.I. [-0.1, 0.01]), where no significant differences were observed (Table 1). To further explore the spatial distribution of this differential effect, we examined its dispersion across the brain networks of interest (Fig. 3c). Our findings indicated a uniform distribution among these networks (X²(3,379) = 0.25, p = 0.8), suggesting that the contribution to this phenomenon is widespread across cortical regions and networks, with no specific network dominance.

**Figure 3:**
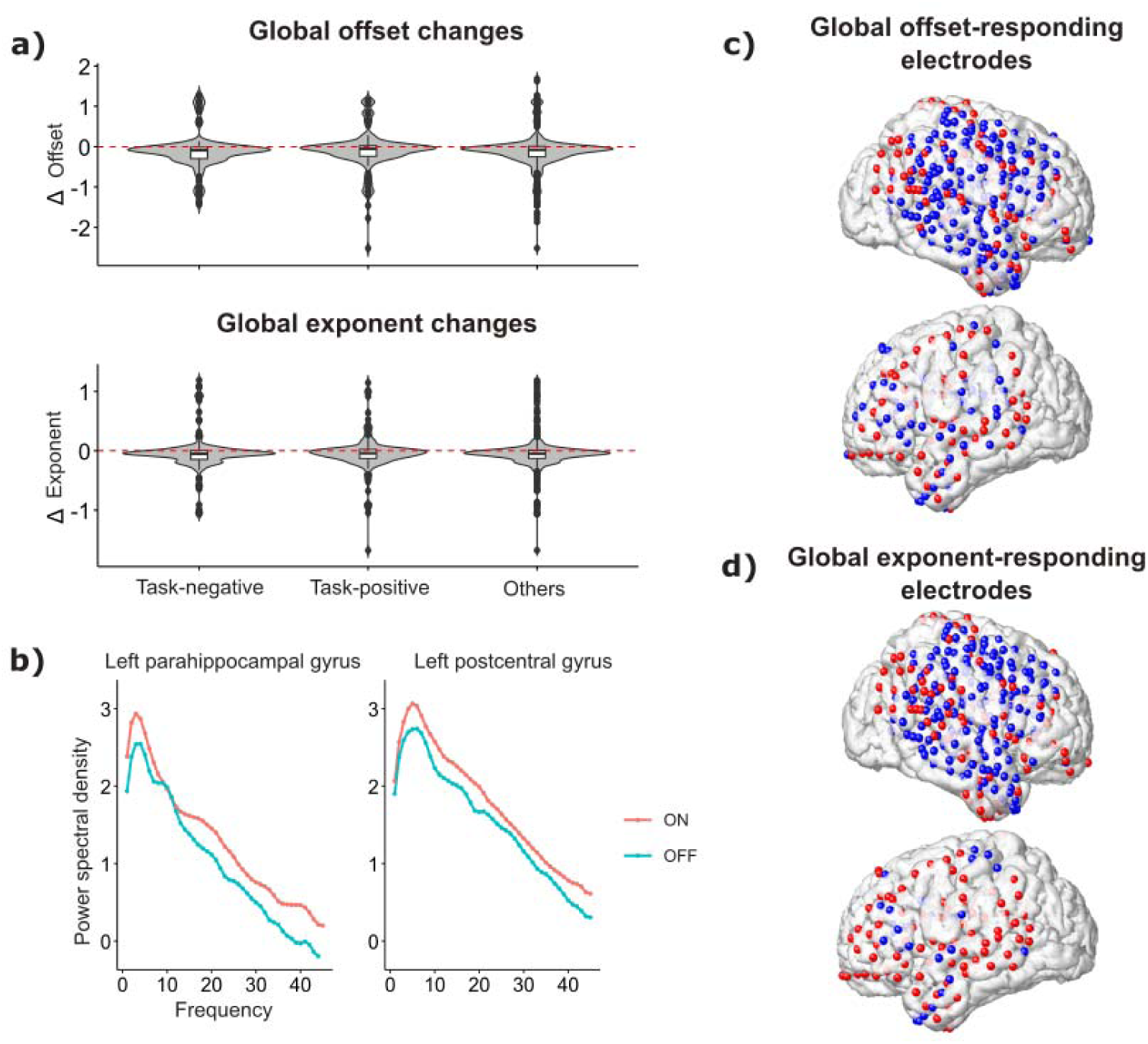
Aperiodic Neural Activity During Mind-wandering. (a) Population distribution of the offset and exponent components of aperiodic activity differences across networks of interest. Negative values indicate a decrease in the aperiodic component during the OFF state. (b) Power spectral density estimates from two spatially distinct electrodes (S1), illustrating the observed aperiodic differences. (c,d) Electrodes contributing to the decrease in offset and exponent. Blue electrodes contribute to the observed differences.

**Table 1:** LMM between ON-OFF states for aperiodic neural activity across networks.

|  | Offset |  |  | Slope |  |  |
| --- | --- | --- | --- | --- | --- | --- |
| | $\beta$ | t | CI | $\beta$ | t | CI |
| TN | -0.11 | -2.7 | [-0.2 -0.03] | -0.04 | -1.14 | [-0.09 0.02] |
| TP | -0.1 | -2.1 | [-0.19 -0.007] | -0.05 | -1.6 | [-0.1 0.01] |
| OTHERS | -0.14 | -3.2 | [-0.2 -0.05] | -0.05 | -2.3 | [-0.1 -0.008] |
| ALL | -0.12 | -4.7 | [-0.1 -0.06] | -0.05 | -2.9 | [-0.08 -0.02] |

$\beta$ Values ( $\beta$ ), t-values (t), and Confidence Intervals (CI) for Task-Positive (TP), Task-Negative (TN), and Other Networks (OTHERS). ALL represent all electrodes.

In conclusion, we observed a generalized decrease in aperiodic components across the brain. Notably, the effects were attenuated when examining individual networks, suggesting a widespread imbalance in inhibitory dominance that may be crucial for the facilitation of MW.

### Global Increases in Connectivity During OFF States

We next evaluated intra-network synchronization in the TN network using the Phase Locking Value (PLV) metric, a widely recognized measure for assessing communication between distant brain regions [26,32,53,54]. Analysis of instantaneous PLVs within the theta band revealed significantly greater phase synchronization in intra-TN networks (β(2,1017) = .01; SE = .002; t = 7.65; p < .01; CI [.01, .02], Permutation test: p < .0001; effect size = .22), with median values of 0.17 ± 0.1 in the ON state and 0.2 ± 0.1 in the OFF state (Fig. 4a, top panel). Similarly, an increase in phase synchronization was observed in intra-TP networks (β(2,943) = .01; SE = .002; t = 4.74; p < .01; CI [.006, .01], Permutation test: p < .0001; effect size = .24), where PLV reached 0.17 ± 0.12 during ON periods compared to 0.21 ± 0.12 in OFF periods (Fig. 4a, middle panel). Finally, intra-OTHER networks also showed significant modulation (β(2,1153) = .004; SE = .001; t = 2.28; p = .02; CI [.0006, .008], Permutation test: p < .001; effect size = .17), with PLV values of 0.17 ± 0.13 and 0.2 ± 0.13 for ON and OFF states, respectively..

**Figure 4:**
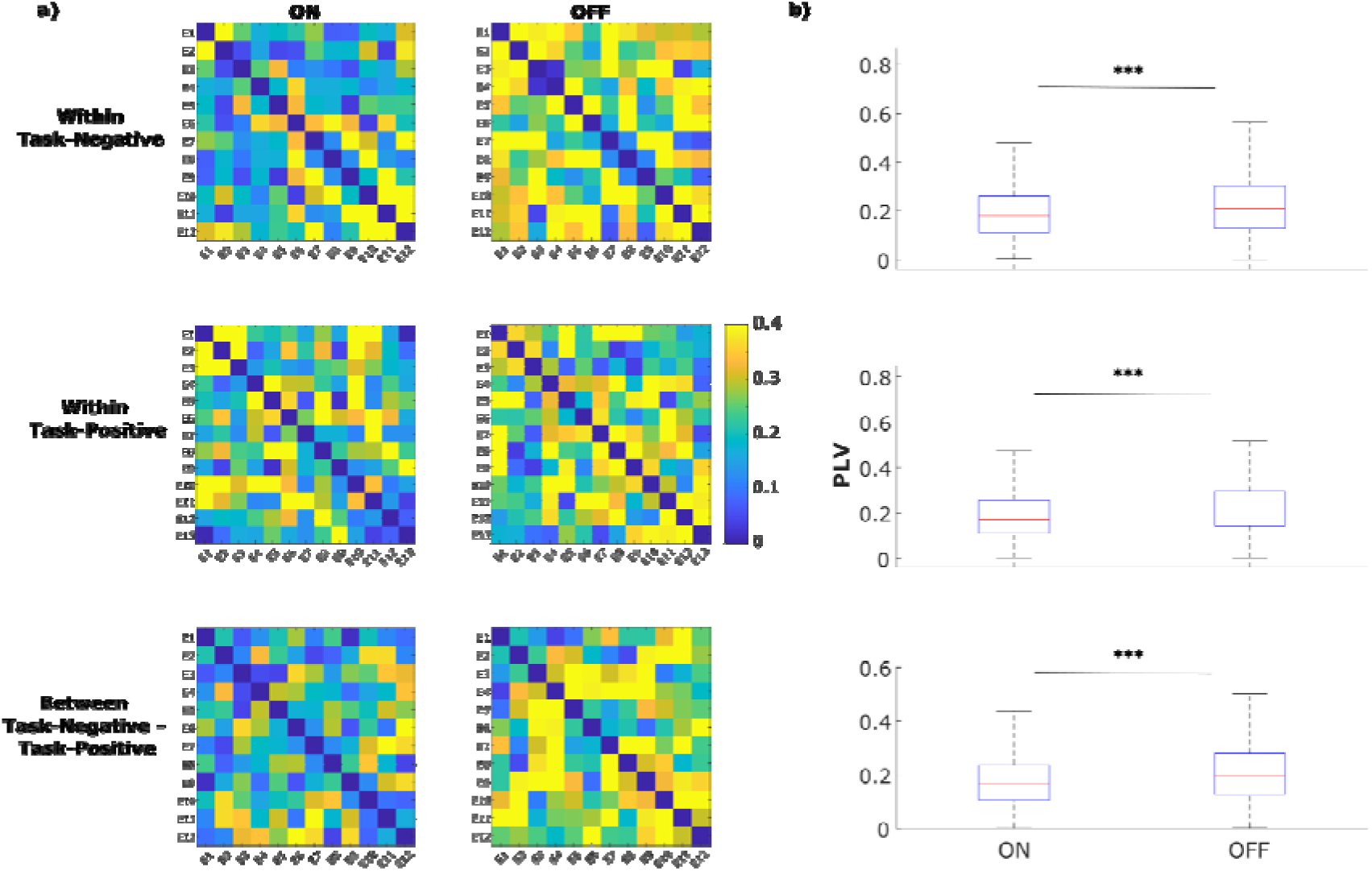
Connectivity Patterns During OFF States. (a) Example connectivity matrix (S1) differentiating between ON-OFF states for: within-task negative connectivity (top), within-task positive connectivity (middle), and between-task negative and task positive connectivity (bottom). (b) Population-level connectivity differences (PLV) corresponding to the matrices shown in a. *** p< .001

**Figure 4.**
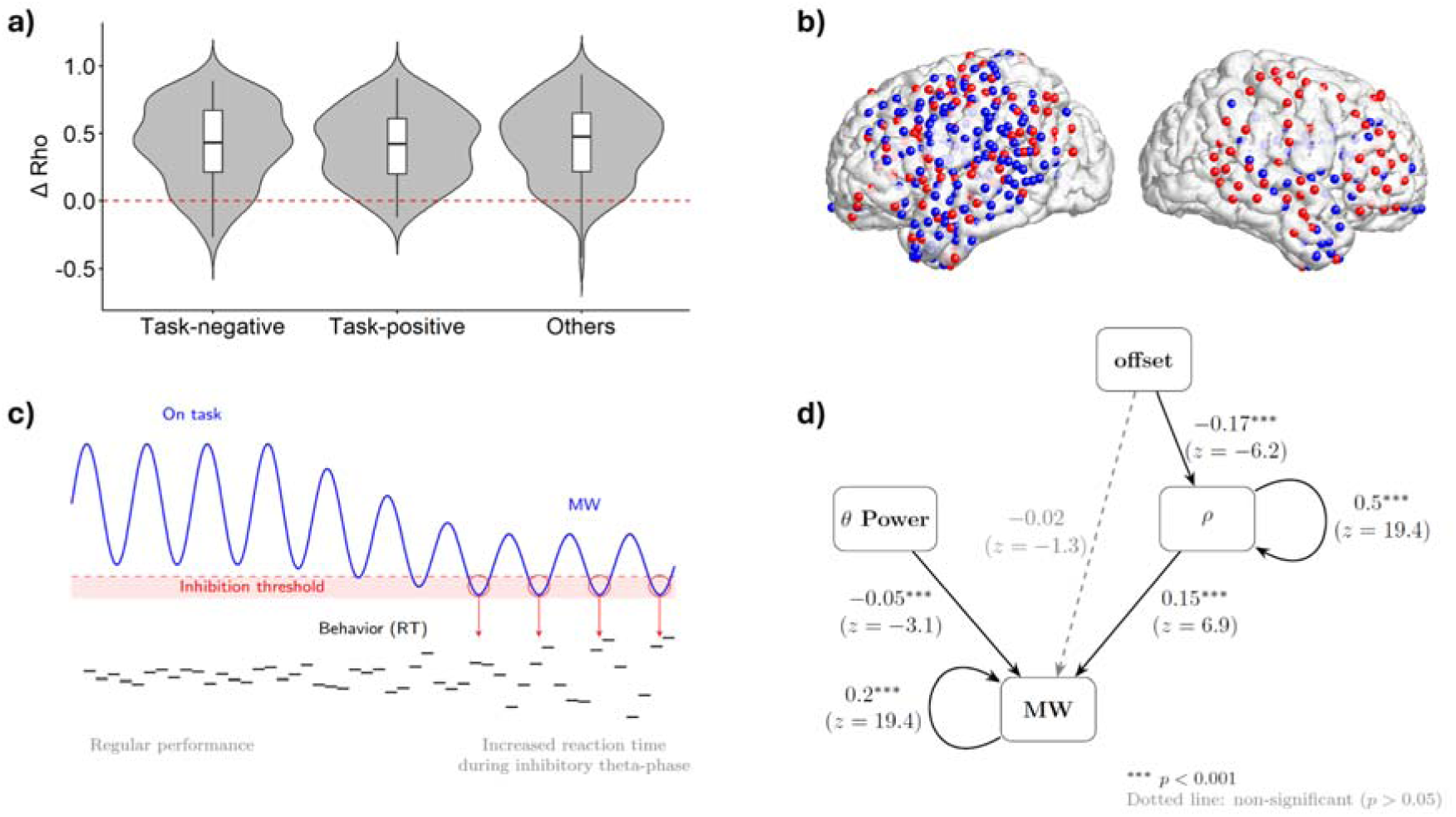
Spontaneous theta dynamics and aperiodic activity jointly relate to mind-wandering. (a) Difference in the circular–linear correlation coefficient (ρ) between theta phase and reaction time (RT) computed as OFF − ON, shown across networks of interest. Positive values indicate stronger phase–behavior coupling during OFF states. (b) Spatial distribution of electrodes contributing to the OFF–ON increase in ρ across cortical networks. (c) Schematic illustration of the proposed mechanism underlying attentional fluctuations. The blue trace represents ongoing theta-band oscillations, and the red shaded segment denotes a transient inhibitory-like window associated with specific theta phases and reduced aperiodic activity. In this framework, these windows coincide with slowed responses (horizontal black segment) and increased trial-to-trial RT variability. (d) Structural equation model (SEM) formalizing the hypothesized relationships between aperiodic offset, theta phase–RT coupling (ρ), and theta power in predicting attentional state (ON vs OFF). Standardized path coefficients (β) and z-values are displayed.

Significant increases in cross-network phase synchronization were also detected between TN-TP (β(2,1900) = .01; SE = .001; t = 7.92; p < .01; CI [.008, .01], Permutation test: p < .001; effect size = .25), with median values of 0.17 ± 0.1 in the ON state and 0.2 ± 0.1 in the OFF state (Fig. 4a, bottom panel). Similar patterns were observed for TN-OTHER (β(2,2206) = .01; SE = .001; t = 10.4; p < .01; CI [.01, .02], Permutation test: p < .001; effect size = .34), where the median PLV was 0.16 ± 0.1 during ON periods compared to 0.18 ± 0.11 in OFF periods. Finally, TP-OTHER connectivity also showed significant modulation (β(2,1017) = .01; SE = .002; t = 7.65; p < .01; CI [.01, .02], Permutation test: p < .001; effect size = .21), with median PLVs reaching 0.2 ± 0.12 and 0.23 ± 0.14 for ON and OFF states, respectively.

Collectively, these findings demonstrate that electrodes within the TN, TP, and OTHER networks exhibit significantly heightened phase synchronization during OFF states. This trend extends to all cross-network interactions, suggesting that the transition to the OFF state is characterized by a widespread increase in functional coupling across the entire recorded ensemble.

### Theta Phase correlated with Attentional State and Behavioral Variability

Thus far, we have observed a generalized decrease in low-frequency bands across brain networks, accompanied by a widespread increase in connectivity at the brain level. Additionally, there is a balancing of inhibitory gains distributed throughout the cortex. The distinct phenomena that differentiate selective attention from MW states are preferentially observed within a particular window of ongoing brain activity, potentially explaining the basis of response variability. Since periods of selective attention are cyclical in nature [39,55] and the oscillatory phase at the onset of each trial is crucial to performance [8,9], we investigated how the instantaneous phase before trial onset relates to subjects’ performance by distinguishing between ON and OFF states for each network of interest. Specifically, for each electrode, we computed a circular-linear correlation between theta phase and RT on a trial-by-trial basis. The resulting ρ values were then analyzed for attentional modulation using an LMM, following the approach of the previous analysis (see Methods for details).

Notably, within the TN network, there is a greater correlation between phase and RT during MW (z = 11.28, p < 0.001). The same pattern is observed in the TP and OTHER networks, where theta phase and RT stability increase in OFF states (z = 11.48, p < 0.01; z = 11.8, p < 0.01, respectively). The Rho difference between OFF and ON states shows a positive values distributed across networks (Fig. 5a), indicating a generalized increase in cortical-level correlation. Electrodes associated with a positive effect on correlation are depicted in Figure 5b.

### The power and phase of theta oscillations together predict the attentional state

Finally, we evaluated the independent contributions of three electrophysiological features in predicting attentional shifts. Using a logistic mixed-effects model to estimate the likelihood of entering a MW state, we further validated the relationship between behavioral and electrophysiological measures. Both theta power (β = −0.27, SE = 0.07, z = −2.85, p < 0.01) and the ρ of the theta phase–RT correlation (β = 0.69, SE = 0.1, z = 6.58, p < 0.01) significantly predicted attentional shifts. In contrast, aperiodic components showed no significant effects (Slope: β = −0.03, SE = 0.15, z = −0.25, p = 0.8; Offset: β = −0.08, SE = 0.14, z = −0.57, p = 0.5). However, the aperiodic offset was significantly correlated with the ρ of phase–RT correlation (β = −0.22, SE = 0.05, t = −4.2, p < 0.001).

Based on the mechanistic model in Fig. 5c, our findings suggest that OFF states are associated with transient theta-phase windows that coincide with slower behavior and increased reaction-time variability. Within this framework, changes in the aperiodic offset may reflect upstream fluctuations in cortical excitability that are compatible with the emergence of these inhibitory-like phases, which in turn may strengthen the circular–linear theta phase–RT coupling (ρ). Theta power, in contrast, may contribute as an additional marker of attentional state, either independently or by modulating the expression of phase-dependent effects. This motivated the use of structural equation modeling to evaluate the hypothesized directional relationships among offset, ρ, theta power, and mind-wandering state. As inputs, we used electrophysiological metrics (theta power, aperiodic offset, and theta phase–RT coupling ρ), controlling for trial count (see Methods), and as output we modeled attentional state (ON vs OFF). Structural equation modeling (SEM; Fig. 5d) supported the hypothesized, including a direct path from offset to state, theta power and ρ were both significantly associated with attentional state (power: β = −0.118, z = −3.15, p = 0.002; ρ: β = 0.246, z = 6.96, p < 0.001), whereas the direct association between aperiodic offset and attentional state was not significant (β = −0.053, z = −1.39, p = 0.165). In contrast, aperiodic offset showed a robust association with ρ (β = −0.222, z = −6.26, p < 0.001), consistent with an indirect relationship between offset and attentional state mediated by phase–RT coupling. The SEM showed excellent fit to the data (CFI = 1.00, RMSEA = 0.00, SRMR = 0.001; χ²(1) = 0.01, p = 0.921).

Overall, these SEM results are consistent with the mechanistic framework proposed in Fig. 5c, in which fluctuations in aperiodic activity are linked to stronger theta phase–behavior coupling (ρ), while theta power and ρ provide complementary markers associated with transitions toward OFF states.

## Discussion

This study investigated the electrophysiological correlates of MW during a sustained attention task. Our findings indicate that global neural oscillations contribute critically to attentional fluctuations and performance variability, extending beyond the influence of localized network activity. Specifically, we identified a widespread reduction in low-frequency power (theta and alpha bands), increased large-scale connectivity, a shift in the excitatory/inhibitory balance, and an increase in the relation between theta phase and reaction time—all of which predict entry into MW in a structured manner.

Our behavioral results reveal that MW is associated with increased RT variability, particularly in the last four trials preceding stimulus presentation. Notably, neither RTs nor accuracy alone reliably distinguished MW from sustained attention, reinforcing the idea that variability in motor responses is a key behavioral marker of MW [48]. These findings are consistent with previous studies suggesting that MW emerges gradually rather than as an abrupt state shift [50,56–58]. The temporal dynamics of MW align with prior work showing that behavioral variability accumulates over time, making multi-trial assessments more predictive of attentional disengagement than discrete error measures [48,50,51]. Moreover, research in perceptual neuroscience has demonstrated that spontaneous neural activity precedes and modulates cognitive and perceptual events, supporting our observation that pre-trial oscillatory activity influences attentional state transitions [1,59,60].

Our electrophysiological findings indicate that MW is characterized by a widespread reduction in theta and alpha power across task-positive (DAN, VAN, FPC) and task-negative (DMN) networks. This non-network-specific suppression challenges models that attribute MW exclusively to default mode network activity [61] and instead suggests a global shift in oscillatory dynamics regulating attentional fluctuations. While EEG studies have reported conflicting results regarding low-frequency oscillations during MW [6,62–64], our intracranial recordings provide a more precise view, revealing consistent low-frequency suppression across cortical networks. These findings align with the hypothesis that MW involves a shift toward internally directed cognition, where sensory and motor engagement is reduced [3,65].

In relation to neural oscillatory activity, the theta band play a critical role in large-scale brain communication, coordinating neural activity across distributed regions [39,66–68]. In our study, theta phase predicted RT variability in MV, suggesting that MW is associated with altered temporal coordination of neural activity. This supports the idea that attentional states are governed by rhythmic neural processes that regulate sensory and motor engagement.

Beyond the periodic component of oscillatory activity, we observed a significant reduction in aperiodic spectral components (exponent and offset) during MW, reflecting a shift in the global excitatory/inhibitory balance. These findings align with previous research showing that decreased aperiodic activity is linked to states of reduced cortical excitability, such as sleep and anesthesia [46,47,69,70]. Aperiodic components provide a key index of neural gain control, reflecting shifts in synaptic activity [44]. Our results suggest that MW may be supported by a global inhibitory state in which cortical excitability is reduced, facilitating a transition away from external task engagement. This interpretation aligns with models proposing that MW serves as a homeostatic mechanism to regulate and respond to cognitive demand [71]. Interestingly, while prior research has emphasized DMN dominance during MW, our results indicate that, in the absence of specific and unique network activity, both periodic and aperiodic activities collectively capture the deployment of MW. This distinction supports the view that MW emerges from a global, inhibitory state that facilitates introspective thought, rather than from activity within a single functional network [72,73].

Brain connectivity plays a crucial role in shaping cognitive states [26,33]. Our findings reveal increased functional connectivity within and across attentional networks during MW, supporting the idea that MW is a distinct neural state rather than a passive disengagement from external tasks. Early connectivity studies using fMRI identified increased DMN coherence during MW [74–76]. However, electrophysiological research has shown that MW is associated with a more complex reorganization of large-scale network interactions [41,77–79]. Our findings suggest that MW is not merely a failure of attention but an active cognitive mode that reorganizes neural communication to support internally generated thought. Importantly, our results highlight that connectivity changes during MW are not restricted to the DMN but extend across task-positive and other networks. This supports a non-hierarchical model of MW, in which attentional fluctuations arise from dynamic interactions across the whole brain rather than from isolated network activity.

Our study provides compelling evidence that MW is characterized by broadly distributed neural oscillations that transcend localized network activity. While the observed reductions in aperiodic slope indicate a shift in the E/I balance, we interpret the “OFF” state primarily as an active reallocation of attentional resources rather than a passive byproduct of global drowsiness. Although these aperiodic shifts mirror patterns seen in anesthesia or deep sleep, we propose that attentional decoupling requires a specific intermediate inhibitory substrate positioned between sustained attention and overt sleep. In this framework, MW may function as a regulatory “windshield wiper” for external cognitive load—a ubiquitous mechanism necessary to mitigate the metabolic demands imposed by continuous stimulus processing. By explicitly excluding sleep trials, we demonstrate that the reported brain connectivity increases and aperiodic activity shifts are intrinsically linked to the active dynamics of the “OFF” state, rather than the physiological signatures of incipient sleep.

These global oscillatory signatures provide a neurophysiological framework that could inform targeted interventions for attentional dysregulation in neuropsychiatric disorders. The identified link between theta phase and behavioral variability suggests potential biomarkers for diagnosing or monitoring conditions such as ADHD, depression, or anxiety, where attentional fluctuations are prevalent [34,80]. Furthermore, the widespread cortical inhibition during MW, reflected in reduced aperiodic components, may represent the mechanism impairing decision-making in clinical populations, potentially informing therapies that target E/I balance. Specifically, neuromodulation techniques such as transcranial magnetic or alternating current stimulation could be tailored to modulate low-frequency activity and enhance attentional control [7,81–83]. Collectively, these insights offer a foundation for personalized therapeutic strategies that leverage neural dynamics to optimize attention and performance across clinical and educational settings.

However, several limitations inherent to invasive neurological protocols must be considered. The imbalance between “ON” (∼510) and “OFF” (∼80) state trials may have reduced the statistical power to detect subtle neural correlates, while asymmetric electrode coverage—with limited representation of the right hemisphere and primary sensory areas—constrains our ability to fully characterize the lateralization of these global dynamics. Additionally, the relatively small cohort size affects generalizability and underscores the need for replication in larger datasets.

Despite these limitations, our findings provide an integrative mechanistic framework—grounded in intracranial electrophysiology, computational modeling, and behavioral analysis—showing that global oscillatory dynamics, rather than passive fluctuations in arousal, shape the architecture of human cognition. Future research should explore how these findings generalize to naturalistic settings and whether interventions targeting these specific oscillations can reliably modulate MW to improve sustained attention.

## Methods

### Subjects

The study cohort comprised nine subjects (S1–S9) undergoing neurosurgical treatment for pharmaco-resistant epilepsy. Data were acquired between 2021 and 2024 at Hospital Dr. Sótero del Río (Puente Alto, Chile) and Hospital Clínico UC Christus (Santiago, Chile). Comprehensive clinical characterization of the cohort—including the anatomical localization of epileptogenic zones and subsequent surgical resection sites—is detailed in Table 2.

**Table 2.** Participant Characteristics: Age, Gender, and Clinical Description of Epilepsy. F: female; M: male; SFE: structural focal epilepsy; FCD: focal cortical dysplasia; MRI: magnetic resonance imaging; PB: phenobarbitone; LEV: levetiracetam; CBZ: carbamazepine; TPM: topiramate; LCM: lacosamide; LTG: lamotrigine; VPA: valproic acid; CLN: clonazepam; CLB: clobazam.

| Patient ID | Age (years) | Gender | Number of records | Diagnosis | Epilepsy duration (years) | Pharmacological treatment | Surgical resection | Seizure onset zone |
| --- | --- | --- | --- | --- | --- | --- | --- | --- |
| S1 | 41 | F | 1 | Left temporal SFE by anti-GAD65 auto antibodies. | 4 | PB, CLB, LEV, LTG, LCM | Polar, lateral, and mesial left temporal resection | Left lateral superior temporal gyrus |
| S2 | 18 | F | 2 | SFE Left precuneus and cingulate gyrus by FCD type IIb | 5 | CLB, VGB, LEV, LCM | Left precuneus and posterior cingulate gyrus | Left precuneus |
| S3 | 16 | F | 3 | SFE Left frontal (lateral F2 and F3 gyrus) by FCD type IIIc (associated with cavernoma) | 3 | CLB, LCM | Medium and inferior lateral left frontal lobe | Medium and inferior lateral left frontal lobe |
| S4 | 28 | F | 2 | SFE right fronto-parietal by perinatal stroke | 28 | LTG, LEV, CLN | Double resection, inferior parietal gyrus, and medium lateral frontal gyrus | Inferior parietal gyrus, and medium lateral frontal gyrus |
| S5 | 25 | F | 2 | Structural focal epilepsy by posterior opercular and insular FCD | 5 | PB, VPA, LEV, LTG, TPM | Triple resection on the left side: inferior lateral frontal gyrus, superior temporal gyrus and temporo-parietal junction | Inferior lateral frontal gyrus, and superior temporal gyrus |
| S6 | 42 | M | 2 | Probable structural focal epilepsy right temporal lobe with negative MRI | 9 | LEV, LTG, TPM | Classic lobectomy of lateral temporal and hippocampectomy on a right side | Right medial temporal lobe |
| S7 | 33 | F | 1 | SFE by left Hippocampal sclerosis | 11 | LEV, LCM | Classic lobectomy of lateral temporal and hippocampectomy on a left side | Left medial temporal lobe |
| S8 | 18 | F | 1 | SFE by fronto-temporoinsular FCD | 8 | CBZ, TPM | Classic lobectomy of lateral temporal and hippocampectomy on the right side plus anterior insula resection | Right lateral temporal and anterior insula |
| S9 | 26 | M | 1 | FCD left posterior temporal lobe | 20 | LEV, LTG, TPM | Left posterior temporal sulcus | Left posterior and lateral temporal sulcus |

Inclusion criteria were: (1) Adult neurology admission; (2) diagnosis of drug-resistant epilepsy according to established clinical criteria; and (3) literacy (ability to read and write). Exclusion criteria included: (1) prior diagnosis of a neurodegenerative disease; (2) presence of developmental disorders; (3) history of significant neurological conditions (e.g., traumatic brain injury); and (4) significant communication impairments.

All participants provided written informed consent prior to participation, including specific consent for the publication of their data. All experimental procedures were approved by the Ethics Committee for Health Sciences at Pontificia Universidad Católica de Chile (Protocol ID: 200811002) and were conducted in accordance with Chilean national regulations, institutional guidelines, and the Declaration of Helsinki.

### Statistical Power Analysis

Based on a previous study that employed the same behavioral task[48], we anticipated a large effect size for the effect of ON and OFF trials on reaction times during the period preceding the question (d = 1.05). A sensitivity power analysis was conducted using G*Power (Version 3.1) to determine the adequacy of our sample size. Given the challenges associated with obtaining intracortical recordings from patients with epilepsy, the sample was necessarily limited by the availability of such rare data. The analysis indicated that a sample size of 8 subjects would be sufficient to achieve an actual power of 0.82 (α = .05). We were able to include 9 subjects in our final sample, which further strengthens the robustness and reproducibility of the findings

### Implants

Subdural electrode arrays (grids and strips; AD-TECH Medical Instrument Corp.) were implanted over one (n = 8) or both hemispheres, with placement guided exclusively by clinical requirements for resective surgery. Electrode contacts were circular (0.8 mm diameter exposed surface) with a 3.5 mm inter-electrode pitch. Following multi-day monitoring, electrodes localized to epileptogenic zones were identified through a multidisciplinary clinical review—including visual inspection of electrocorticography (ECoG) recordings, seizure onset localization, and functional mapping of eloquent cortex—and subsequently excluded from the analysis.

### Experimental Task

The Sustained Attention to Response Task (SART) was administered using a protocol adapted from our prior research [48] . The task was presented at the bedside to each patient across multiple sessions (range: 1–3). Each session consisted of three to nine blocks, with a maximum of three blocks administered per day. Each block lasted approximately nine minutes. To ensure consistency across clinical sites, the task was delivered at the bedside using the same laptop model connected to an external 21-inch monitor positioned 60 cm from the patient at chest level. Stimulus presentation was controlled using Presentation® software (Version 23.0, Neurobehavioral Systems, Inc., Berkeley, CA).

During the SART, participants viewed a rapid serial visual presentation of ten randomly selected uppercase letters against a gray background. Each letter was displayed for 100 ms inside a central white rectangle, followed by a variable inter-stimulus interval (ISI) ranging randomly from 2,500 to 4,000 ms. In this framework, a single trial was defined as the duration encompassing the stimulus onset (either target or non-target) and the subsequent ISI. Participants were instructed to press one button for the infrequent target letter “F” (occurring in 10% of trials) and a different button for all other non-target letters. This design establishes a prepotent “go” response, requiring participants to sustain attention to correctly withhold responses on rare target trials.

To probe fluctuations in subjective attentional state, Thought Sampling Questions (TSQs) were pseudorandomly presented within each block, occurring every 5 to 8 trials. Each TSQ posed the statement: “Just before this question, your attention was distracted by:” accompanied by three response options: (1) “My attention was not distracted,” (2) “Losing vigilance (falling asleep),” and (3) “Internal distraction (memories, imagination, thoughts).” Participants responded using a numeric keypad. These responses allowed us to classify moments of task engagement into three mental states: an “ON” state of sustained attention (Response 1), an “OFF” state of mind-wandering (Response 3), and an “SL” state indicative of drowsiness or sleep onset (Response 2).

Across all participants, a total of 646 TSQ responses were collected. The distribution of states was as follows: 510 “ON” responses (78.9%), 80 “OFF” responses (12.4%), and 56 “SL” responses (8.7%). Notably, four participants reported no “SL” responses, further underscoring individual variability in self-reported task engagement.

### Electrocorticography Data Acquisition

ECoG recordings were acquired at the bedside within the subjects’ clinical suites. Data were recorded using a Cadwell Easy III system (Cadwell® Industries Inc.) at a sampling rate of 250 Hz with an integrated 50 Hz notch filter, as well as via a Natus Quantum 128-channel system (Natus® Medical Inc.) sampled at 1024 Hz. The spatial distribution and configuration of the implants—including total electrode counts and the final subset of viable electrodes per subject—are summarized in Table 3.

**Table 3.** Implant Characteristics: Type of Implant, Number of Functional Electrodes, and Number of Electrodes in Networks of Interest for Each Patient. DMN: default mode network; VAN: ventral attention network; DAN: dorsal attention network; SM: sensorimotor network; LIM: limbic network.

| Patient | Implants | Sampling rate (Hz) | # total electrodes | # epileptogenic electrodes | # useful electrodes | # electrodes within networks |
| --- | --- | --- | --- | --- | --- | --- |
| S1 | 2 strips<br>2 grids | 250 | 62 | 14 | 48 | DMN: 12<br>VAN: 4<br>DAN: 10<br>SM: 11<br>LIM: 11 |
| S2 | 2 strips<br>1 grid | 250 | 40 | 11 | 29 | DMN: 12<br>VAN: 2<br>DAN: 1<br>SM: 9<br>LIM: 0 |
| S3 | 1 grid | 250 | 64 | 18 | 26 | DMN: 4<br>VAN: 2<br>DAN: 4<br>SM: 9<br>LIM: 0 |
| S4 | 1 grid<br>1 strip<br>1 deep electrode | 250 | 74 | 12 | 41 | DMN: 13<br>VAN: 2<br>DAN: 6<br>SM: 9<br>LIM: 0 |
| S5 | 1 grid | 1024 | 64 | 10 | 54 | DMN: 20<br>VAN: 7<br>DAN: 4<br>SM: 17<br>LIM: 2 |
| S6 | 3 strips<br>1 grid | 250 | 84 | 5 | 56 | DMN: 23<br>VAN: 4<br>DAN: 7<br>SM: 13<br>LIM: 7 |
| S7 | 2 strips | 1024 | 12 | 6 | 6 | DMN: 2<br>VAN: 0<br>DAN: 0<br>SM: 0<br>LIM: 4 |
| S8 | 2 strips<br>1 grid | 250 | 78 | 6 | 55 | DMN: 16<br>VAN: 4<br>DAN: 3<br>SM: 10<br>LIM: 5 |
| S9 | 1 strip<br>1 grid | 1024 | 72 | 4 | 68 | DMN: 24<br>VAN: 5<br>DAN: 10<br>SM: 15<br>LIM: 10 |

### ECoG data preprocessing

Continuous ECoG recordings were segmented into epochs strictly corresponding to the task duration. A notch filter was applied to attenuate power-line noise, with a band-stop range of 48–52 Hz.

Signals from each channel were initially re-referenced to the common average of all electrodes within the same strip or grid. For connectivity analyses, however, we applied a current source density (CSD) transformation, in which each electrode was re-referenced to the weighted average of its neighboring electrodes (typically 2–4 neighbors, depending on grid position). This approach enhances spatial specificity by attenuating globally shared signals and reducing the impact of volume conduction [84,85]. Channels were excluded from the average if they exhibited pathological activity during clinical monitoring (as determined by a neurologist), had a variance exceeding or falling below five times the median variance across all channels, or displayed more than three times the median number of spikes, defined as abrupt 100 μV changes between successive samples. In addition, the data were sub-sampled at 250 Hz to homogenize sampling frequencies between devices.

We then segmented the data into time windows aligned to the onset of letter presentation (trial onset). Each 3.5-second window ranged from -2 s to +1.5 s relative to trial onset and represented the average activity across the last four trials preceding the TSQ, ensuring consistency with the behavioral results (Suplementary Figure 1). We analyzed the 1–90 Hz frequency band for both pre- and post-stimulus periods. Time-frequency representations were computed using a sliding Hanning window of 400 ms duration. The window was advanced in 10 ms steps, resulting in a 97.5% overlap between successive temporal bins. Time–frequency (TF) power was computed at the single-trial level for each electrode, time point, and frequency bin. To evaluate the effect of attentional state across the cohort, we employed a linear mixed-effects model (LMM) for each TF bin. The model included attentional state (ON vs. OFF) as a fixed effect. To account for the hierarchical nature of the data, we specified random intercepts and slopes at the electrode level, nested within subjects (e|s), thereby capturing inter-electrode and inter-subject variability.

The model is formalized as:

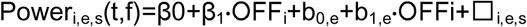

where Power_i,e,s_(t,f) represents the power at time t and frequency f for trial i, electrode e, and subject s; OFF_i_ is a binary variable indicating attentional state (ON = 0, OFF = 1); β_0_ and β_1_ are fixed effects; b_0,e_ and b_1,e_ are random intercepts and slopes at the electrode level (nested within subjects); and □_i,e,s_ represent the gaussian residual error.

Instead, all trial-level observations contributed to the estimation of model parameters. For each TF bin, the t-statistic associated with the attentional state regressor was extracted and used to generate time–frequency statistical maps. To correct for multiple comparisons, we applied false discovery rate (FDR) correction (q < 0.05) across all time × frequency bins.

Note that time–frequency analyses were conducted within a single continuous time window spanning both pre- and post-stimulus periods. Importantly, no explicit baseline normalization (e.g., subtraction or division by a pre-stimulus interval) was applied. Instead, normalization was achieved within the statistical modeling framework. Specifically, the intercept captures the mean power level for each electrode, and fixed effects estimate condition-related deviations from this mean. This approach allows for direct comparison of pre- and post-stimulus activity without introducing biases associated with arbitrary baseline selection. Furthermore, statistical maps were expressed as t-values (β / SE), providing an implicit standardization across frequencies and time points, and mitigating differences in absolute power (e.g., due to the 1/f structure of electrophysiological signals). This framework enables a unified and unbiased characterization of ongoing and evoked dynamics within the same model. For further implementation of such a process see prior work[26,54]. The complete preprocessing and analysis pipeline is summarized in Supplementary Figure 3.

All preprocessing and analyses were conducted using MATLAB (Version 9.8.0, R2020a) and the FieldTrip toolbox (http://fieldtriptoolbox.org).

### Anatomical Localization and Classification of Electrode Contacts

We developed a comprehensive image-processing pipeline to precisely localize ECoG electrode contacts within the brain[86]. This pipeline integrates electrode contact localization, brain tissue segmentation, and alignment with an anatomical atlas, leveraging pre-surgical MRI (T1-weighted, 1.5T) and post-surgical computed tomography (CT) scans.

The process began with the preprocessing and analysis of T1-weighted MRI volumes using the Brainstorm Toolbox[87]. Cortical surface extraction was subsequently performed using CAT12 (https://neuro-jena.github.io/cat/). Pre-operative T1 volumes were standardized to the Montreal Neurological Institute (MNI 152) template at a 1-mm resolution using nonlinear registration according to Jenkinson and Smith’s approach [88]. The obtained normalization matrices were then used to align the registered post-operative volumes with the MNI template.

Co-registration of CT and T1 scans was performed using SPM12, followed by segmentation to reconstruct cortical surfaces. The anatomical labeling of electrodes was based on individualized cortical images co-registered with the CT scans. Individual electrode localization is shown in Supplementary Figure 2. Each electrode location was precisely identified, including tissue probability and anatomical region, by mapping electrode coordinates to the brain tissue and anatomical atlas information.

Finally, electrodes were classified into one of three groups of networks—Task-positive (VAN, DAN, FPC), Task-negative (DMN), or Others (Sensorimotor (SM), LIMBIC) — using the Schaefer atlas of seven cortical networks with 100 parcellations [89]. To achieve this, we registered the Schaefer atlas from fsaverage space to individual cortical surfaces via the Brainstorm Toolbox. Each vertex on the pial surface was then assigned to one of the seven networks, ensuring accurate classification of electrode memberships.

### Connectivity analysis

Connectivity analysis was performed using the Phase Locking Value (PLV) metric [53,90]. Frequency bands of interest (FOI) were defined as Theta (4–8 Hz), and the time window of interest (TOI) was set as the 400 ms preceding trial onset (−400 ms to −10 ms). Electrodes within each network of interest (Task-positive, Task-negative, Others) were selected for analysis. For each electrode within a subject, imaginary Hilbert transform values were computed based on the defined FOI. Subsequently, the PLV was calculated as the absolute average of the angular differences during the TOI for each pair of electrodes. Mathematically [91], the PLV can be expressed as:

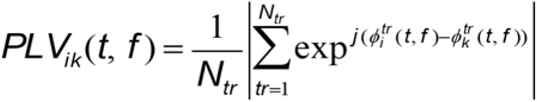

where (Φ) is the phase of a pair of electrodes (i, k) at each time point of interest (t), and (f) represents the total number of frequency points of interest analyzed; (tr) represents the analysis for each trial. This calculation provides a measure of the synchrony between the activities of different electrode pairs within the specified frequency bands and time intervals, thereby enabling insights into the connectivity dynamics among the specified brain networks.

Given the lower number of “OFF” trials in comparison with “ON” trials, we implemented a trial-matching procedure to ensure a balanced and unbiased comparison between conditions. For each “OFF” trial, we selected a corresponding “ON” trial defined as the immediately following trial occurring after the TSQ. This approach ensured homogeneous sampling across the task timeline, preserved the temporal structure of the experiment, and controlled for potential confounds such as order effects or fatigue, while maintaining a clear separation between cognitive states.

To validate that the observed connectivity increases in the “OFF” state were not artifacts of the smaller trial count or signal variability, we performed a permutation test. Condition labels (ON/OFF) were randomly shuffled across trials while preserving the matching structure described above. This procedure was repeated 10,000 times to generate a null distribution of connectivity differences. The observed connectivity differences were compared against this null distribution and considered significant if they exceeded the 95th percentile (p < 0.05).

### Aperiodic Component Analysis

To characterize the aperiodic and periodic features of the neural power spectra, we applied the FOOOF algorithm (SpecParam [42,43]) as implemented in the FieldTrip toolbox (ft_freqanalysis). The analysis was performed on preprocessed time-series data, specifically focusing on the pre-stimulus baseline period (−2 to 0 s) to capture spontaneous oscillatory activity.

Prior to the parameterization, power spectral density (PSD) estimates were computed using a multi-taper frequency transformation (mtmfft) with Discrete Prolate Spheroidal Sequences (DPSS) tapers. We applied a frequency smoothing of 2 Hz (cfg.tapsmofrq = 2) to ensure robust spectral estimation. The algorithm was fitted across a frequency range of [1, 90] Hz using a ’fixed’ aperiodic model (linear fit in log-log space).

Consistent with established best practices, we utilized the default heuristic parameters for peak detection to avoid overfitting: the peak width limits were set to [0.5, 12] Hz, the maximum number of peaks was not explicitly constrained, and a peak threshold of 2 standard deviations was applied. This approach allowed for a clear separation of the 1/f aperiodic background from the physiological oscillations.

### Phase Calculation

The instantaneous phase was derived from the Hilbert time-frequency transform. For this analysis, FOI was Theta (4–8 Hz), and TOI was set to 10 ms before trial onset. Initially, the analytic Hilbert transform was averaged across ON and OFF trials, followed by averaging the imaginary part of the transform.

## Statistical Analysis

Statistical analysis of behavioral data proceeded in three stages. First, target versus non-target trial performance was compared for both hit rates and mean reaction times (RTs) using non-parametric Wilcoxon signed-rank tests. Second, to capture trial-to-trial performance variability, we derived two dynamic RT metrics: the absolute value of consecutive RT differences (|RT□₋₁ − RT□|) and the standard deviation of these differences. Group-level comparisons of these metrics (mean RT, RT difference, and RT variability) preceding specific events (TSQs) were conducted using Kruskal-Wallis tests; significant effects were followed by Bonferroni-corrected Wilcoxon post-hoc tests. Third, we assessed the strength of evidence for condition-dependent effects (ON vs. OFF) using Bayesian regression models, implemented in JASP (Version 0.18.2) with 5,000 iterations.

The statistical analysis of connectivity and aperiodic components was performed using linear mixed-effects models (LMM), with attentional state (ON vs OFF) included as a fixed effect. For these analyses, connectivity metrics (PLV) and aperiodic parameters (slope and offset) were first computed at the electrode (or electrode-pair) level separately for ON and OFF conditions. These condition-specific values were then entered into the LMM. Random effects included electrodes (or electrode pairs) nested within subjects, allowing us to account for variability across recording sites and participants. In contrast, time–frequency power analyses were conducted at the single-trial level (see above), where all trials were included in the model without prior averaging. To visualize spatial effects, we used the sum of the fixed intercept and the fixed effect associated with attentional state as a thresholding criterion. Electrodes (or electrode pairs) showing values below this threshold were plotted in white, whereas those exceeding the threshold were plotted in blue.

The circular-linear correlation coefficient was computed between Theta phase data and RT [92]. Comparisons of Rho (ρ) coefficients between ON - OFF states were made using an LMM as described above. To predict the occurrence of OFF states, we fitted logistic mixed-effects models with electrode-level random intercepts and slopes. We conducted a structural equation model using the lavaan package in R. For both models, all brain metrics were z-scored. To control for potential confounds from variable trial counts for conditions, we regressed out the number of trials using a generalized linear model (GLM) before analysis. This ensured that observed effects were not driven by differences in trial sampling.

To further evaluate the hierarchical relationships between oscillatory power, aperiodic activity, and phase–behavior coupling in predicting attentional state, we implemented a structural equation model (SEM) using the lavaan package in R. Prior to SEM estimation, electrophysiological variables were standardized (z-scored) to enable direct comparison of path coefficients across measures. Because the number of available trials differed across attentional conditions and could introduce systematic variance in these metrics, we controlled for trial-count effects by regressing each standardized metric against the standardized trial count using generalized linear models, and subsequently extracted the residualized variables for SEM analyses. The SEM tested a directed path model consistent with the mechanism illustrated in Fig. 4d, in which mind-wandering state (state, binary outcome) was predicted by residualized theta power and residualized phase–RT coupling (state ∼ theta + ρ), while phase–RT coupling was predicted by residualized aperiodic offset (ρ ∼ offset). This specification allowed us to evaluate both the direct contributions of theta power and phase coupling to attentional state, and the indirect influence of aperiodic offset mediated through ρ. Model parameters were estimated using maximum likelihood with robust standard errors, and overall fit was evaluated using conventional indices including χ², RMSEA, CFI, and SRMR, with thresholds aligned with conventional standards (e.g., CFI > 0.90). We additionally compared this hypothesized model against an alternative model that included a direct path from aperiodic offset to attentional state (state ∼ thea + ρ + offset) using information criteria (AIC/BIC) and likelihood-ratio testing for nested models.

Both behavioral and electrophysiological statistical analysis was performed with R (v 4.2.2), RStudio (v 2023.06.0), and MATLAB (Version 9.8.0, R2020a). The significant level was set to 0.05.

## Data Availability

The complete raw data set underlying the results used in our study will be available in the public repository on github (https://github.com/neurocics/).

## Code Availability

The additional toolbox and codes used in the analysis will be available on our lab github site as LANtoolbox (https://github.com/neurocics/LAN_current) and paper’s name, once accepted for publication .

## Author Contributions

JH, RH-C, RU-SM, PF, FA conceptualized the study. JH, RH-C, RU-SM, CC, PM, and JG collected the data. JH, AF-V, and PB analyzed the data. JH, AF-V, RH-C, PB, and FA wrote the main manuscripts and prepared the figures. All authors reviewed the manuscript.

## Acknowledgments

We sincerely thank the National Reference Center of Hospital Sótero del Río in Santiago, Chile, and the Epilepsy Unit of the Department of Neurology at the Faculty of Medicine of the Pontifical Catholic University of Chile for their support in this study. We are especially grateful to all the patients and their families for their invaluable contribution to this research.

## Supplementary Material

**Supplementary Figure 1.**
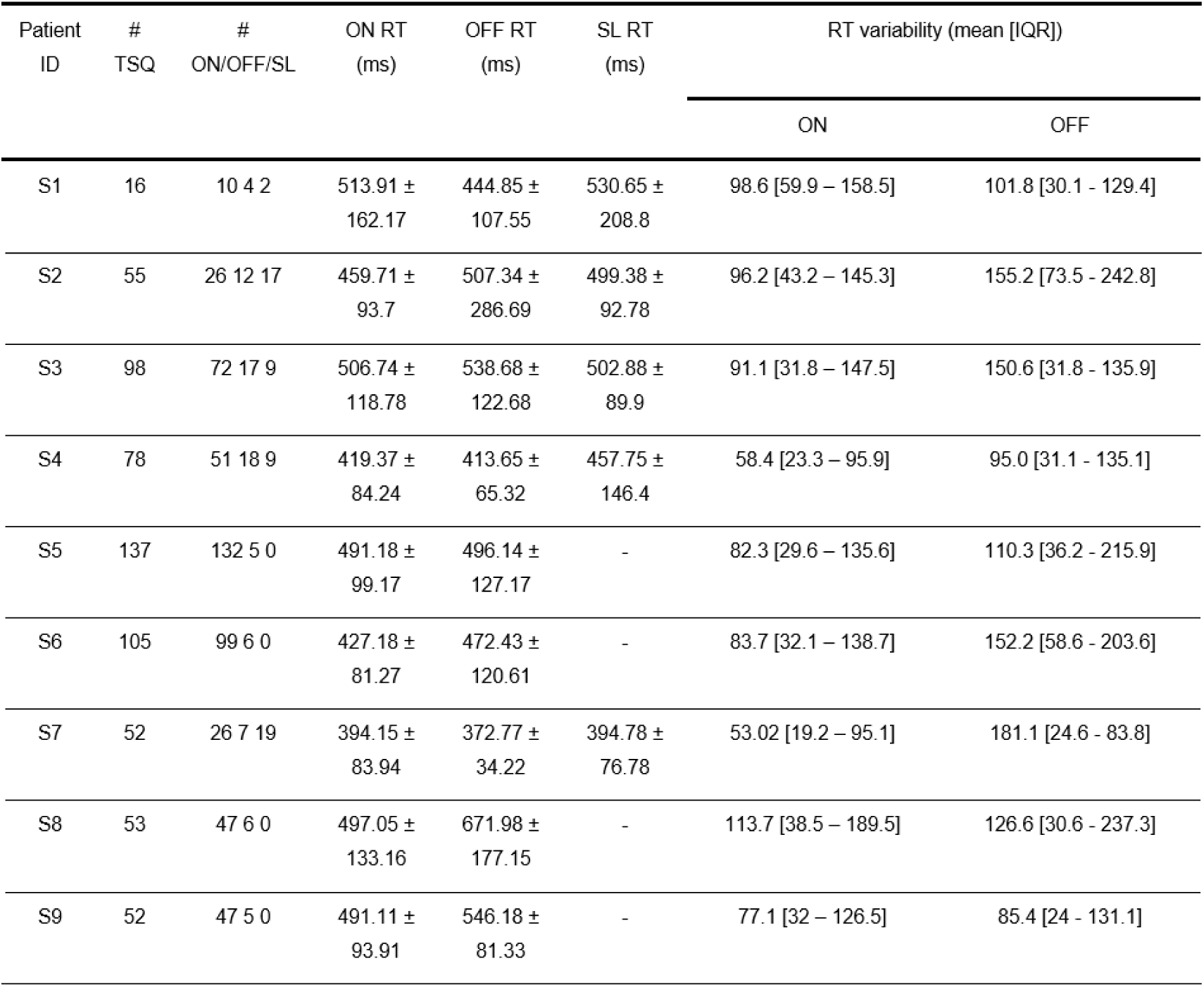
Participant-level descriptive statistics. TSQ = Through sampling question; ms = millisecond; ON = Sustain attention; OFF = mind wandering; SL = Sleep condition; RT = reaction time; IQR: interquartile range. RT are shown as mean ± standard deviation.

**Supplementary Figure 2:**
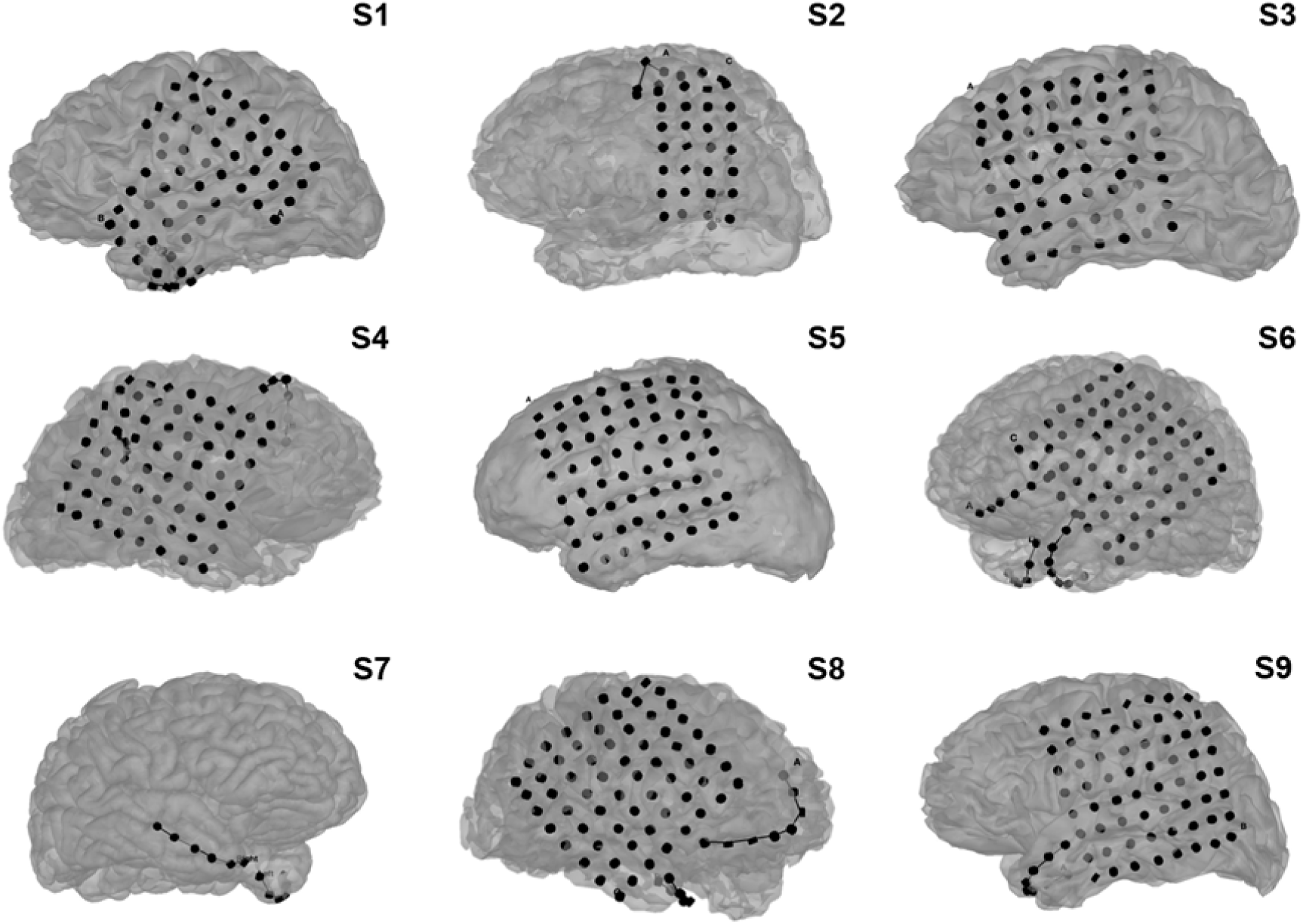
Independent brain electrodes by subject.

**Supplementary Figure 3:**
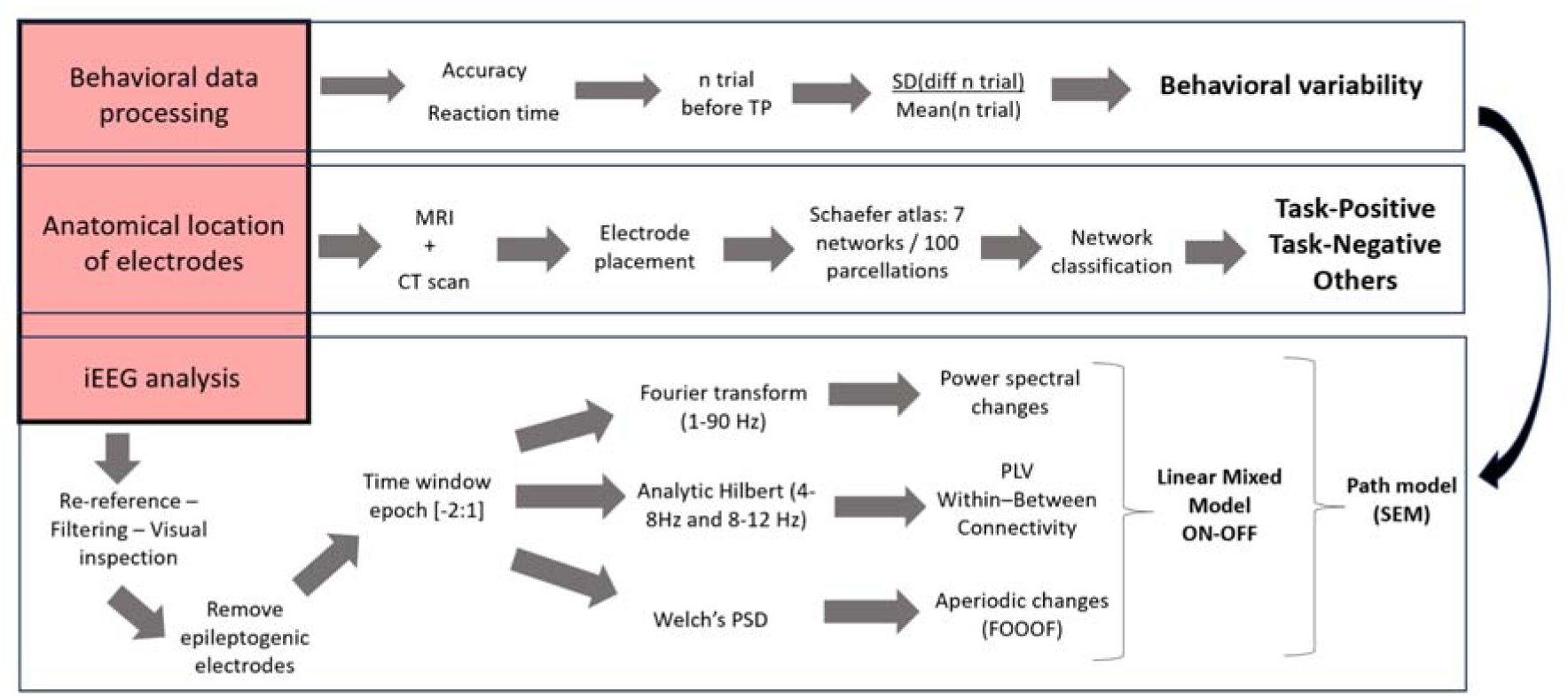
Flowchart of the ECoG signal processing pipeline and electrode-level statistical modeling.

**Supplementary table 1:**
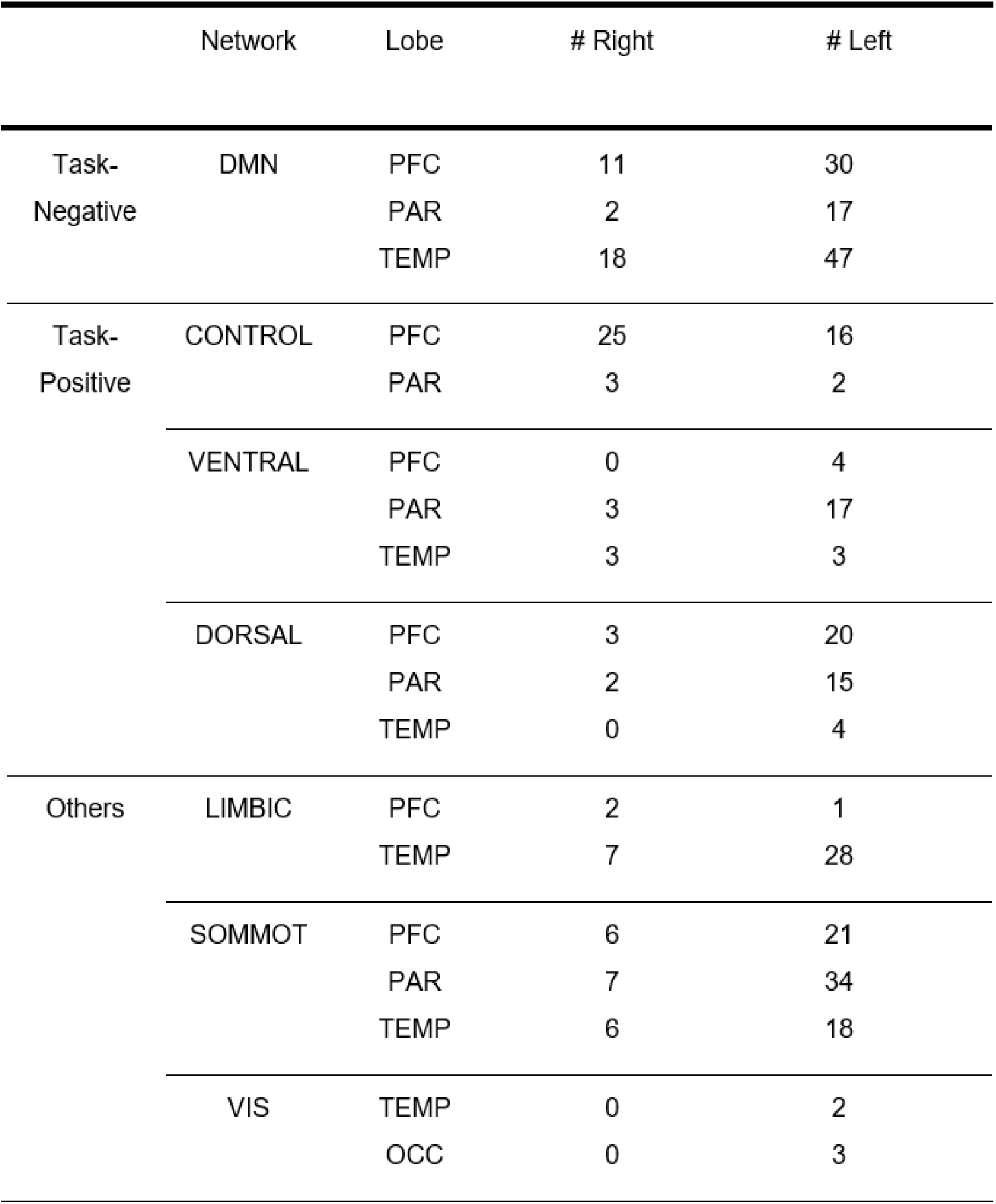
Anatomical electrode distributions.

